# Time-resolved transcriptome of barley anthers and meiocytes reveals robust and largely stable gene expression changes at meiosis entry

**DOI:** 10.1101/2020.04.20.051425

**Authors:** Abdellah Barakate, Jamie Orr, Miriam Schreiber, Isabelle Colas, Dominika Lewandowska, Nicola McCallum, Malcolm Macaulay, Jenny Morris, Mikel Arrieta, Pete E. Hedley, Luke Ramsay, Robbie Waugh

## Abstract

In flowering plants, successful germinal cell development and meiotic recombination depend upon a combination of environmental and genetic factors. To gain insights into this specialised reproductive development programme we used short- and long-read RNA-sequencing (RNA-seq) to study the temporal dynamics of transcript abundance in immuno-cytologically staged barley (*Hordeum vulgare*) anthers and meiocytes. We show that the most significant transcriptional changes occur at the transition from pre-meiosis to leptotene–zygotene, which is followed by largely stable transcript abundance throughout prophase I. Our analysis reveals that the developing anthers and meiocytes are enriched in long non-coding RNAs (lncRNAs) and that entry to meiosis is characterized by their robust and significant down regulation. Intriguingly, only 24% of a collection of putative meiotic gene orthologues showed differential transcript abundance in at least one stage or tissue comparison. Changes in the abundance of numerous transcription factors, representatives of the small RNA processing machinery, and post-translational modification pathways highlight the complexity of the regulatory networks involved. These developmental, time-resolved, and dynamic transcriptomes increase our understanding of anther and meiocyte development and will help guide future research.

**One sentence summary:** Analysis of RNA-seq data from meiotically staged barley anthers and meiocytes highlights the role of lncRNAs within a complex network of transcriptional and post-transcriptional regulation accompanied by a hiatus in differential gene expression during prophase I.

The author responsible for distribution of materials integral to the findings presented in this article in accordance with the policy described in the Instructions for Authors (www.plantcell.org) is: Robbie Waugh

## INTRODUCTION

Along with global population growth and climate change, food security is a major challenge that can be partly addressed by breeding new crop traits. Breeding relies almost entirely upon meiosis during which new allelic combinations are created and segregated by the meiotic recombination machinery. In the life cycle of flowering plants, this pivotal event is induced by developmental and environmental cues and occurs within the specialised reproductive male (stamens) and female (carpel) organs. These are almost entirely composed of vegetative tissues embedding a small number of archesporial cells that differentiate into meiocytes (Goldberg *et al.*, 1993; Hord and Ma, 2007; Kelliher and Walbot, 2011). Understanding the development of these organs and the underpinning biology of meiotic recombination could help improve the efficiency of trait mapping and lead to improvements in crop breeding.

Meiosis is a fundamental eukaryotic process that generates haploid gametes from the parental diploid meiocytes. During this specialised cell division, the parental chromosomes (homologues) condense, pair, synapse, and undergo recombination within a meiosis-specific protein complex called the synaptonemal complex (SC) before they separate in meiosis I (Naranjo, 2012; Osman *et al.*, 2011; Zickler and Kleckner, 2015). The mechanisms which drive the homology search that establishes proper pairing of homologues are poorly understood. In most studied species, the formation of the SC has been shown to be intricately connected to the introduction of programmed DNA double-strand breaks (DSBs) and their repair by the recombination machinery (Naranjo, 2012; Zickler and Kleckner, 2015). The resulting gene shuffling, key to breeding desired traits, happens mainly at the ends of chromosomes while the genes in the middle rarely recombine (Künzel and Waugh, 2002). This bias in the distribution of recombination events is particularly pronounced in the large grass family (Poaceae), including cereal crops (Higgins *et al.*, 2014). Meiotic function is evolutionarily well conserved even though the amino acid sequences of recombination and particularly SC proteins are variable in different species (Grishaeva and Bogdanov, 2017; Wang and Copenhaver, 2018). Phylogenomic, mutagenic, and immuno-cytological tools have been extensively used to clone and characterise plant orthologues of meiotic genes (Lambing *et al.*, 2017).

During early anther development, meristematic cells divide and organise into four pollen sacs where the archesporial cells divide mitotically to generate the primary sporogenous cells surrounded by four concentric tissues mostly made of a single cell layer. The primary sporogenous cells divide further to generate a diploid pre-meiotic sporogenous tissue within each pollen sac (Goldberg *et al.*, 1993). This complex developmental program involves multiple signalling pathways controlled by the cooperation of several receptor-like protein kinases (Cui *et al.*, 2018). The anther is thus a sophisticated organ made of multiple somatic tissues that harbour and support meiotic cells to generate and release the viable pollen that are necessary for successful fertilization and new seed formation. At the early stages, the switch from pre-meiotic germinal cells at the centre of anther lobes to meiotic cells is triggered by changes in gene expression. Using laser microdissection, microarray expression profiling, and mass spectrometry, Yuan *et al.* (2018) conducted analysis of pre-meiotic germinal cell development within maize anthers, and demonstrated that many meiotic genes are both transcribed and translated prior to entry into meiotic prophase I. Mirroring Yuan *et al.* (2018), Nelms and Walbot (2019) investigated transcriptional dynamics in maize anther meiocytes at the point of entry into prophase I using single-cell RNA sequencing, identifying a further two-step transcriptional reorganisation at leptotene. Analysis of the meiocyte transcriptome of domesticated and wild sunflowers revealed a potential regulatory role for long non-coding RNAs (lncRNAs) in meiotic recombination (Flórez-Zapata *et al.*, 2016).

To gain further insight into transcriptional and post-transcriptional regulation in barley reproductive tissues we combined precise immuno-cytological staging with RNA sequencing to analyse transcript profiles of barley anther and meiocyte tissues before, during, and after prophase I. Our data reveal extensive transcriptional reorganisation between pre-meiosis and leptotene which is rapidly stabilised and then persists throughout prophase I. Further, differential and co-expression analyses indicate the potential involvement of several regulatory pathway components in this reorganisation, including numerous transcription factors, RNA binding proteins, E3 ubiquitin ligases, methyltransferases, and lncRNAs. The details captured in our transcript abundance datasets provide significant insights into this highly organised, stepwise biological process and will lead to both greater understanding and possible applied outcomes.

## RESULTS

### Accurate staging of anthers

Meiotic prophase I in barley progresses over an approximate 36-hour time period inside the anthers of developing spikelets (Higgins *et al.*, 2012; Barakate *et al.*, 2014), that are themselves buried within the extending barley culm. To collect appropriately staged meiotic inflorescences, or spikes, we first developed a reliable prediction system based on external morphological characteristics of the main tiller as described for the cultivar Optic (Gómez and Wilson, 2012). In our environmentally conditioned growth room (see materials and methods), the barley cultivar Golden Promise plants reached meiosis at six weeks post-germination, when the flag leaf was emerging, and the spikes were 0.5 to 2 cm in length. As we had previously determined a strong correlation between anther length and meiotic progression (Arrieta *et al.*, 2020), anthers were dissected under a stereomicroscope and measured before being stored as length classes in RNA later™. For each length class, a subset of 10 anthers was stored in PBS buffer and used for chromosomal acetocarmine staining for meiotic staging. This allowed us to determine the precise correlation between anther size and meiotic stage in this material. As meiosis is not completely synchronised along the length of a spike (Lindgren *et al.*, 1969), only anthers from the middle spikelets were collected. In total, three biological replicates with at least 120 anthers in each were collected for pre-meiosis (0.3–0.4 mm anthers), leptotene–zygotene (0.5–0.9 mm anthers), pachytene–diplotene (1.0–1.2 mm anthers) and metaphase I–tetrad (1.3–1.4 mm anthers) (Figure 1).

**Figure 1:**
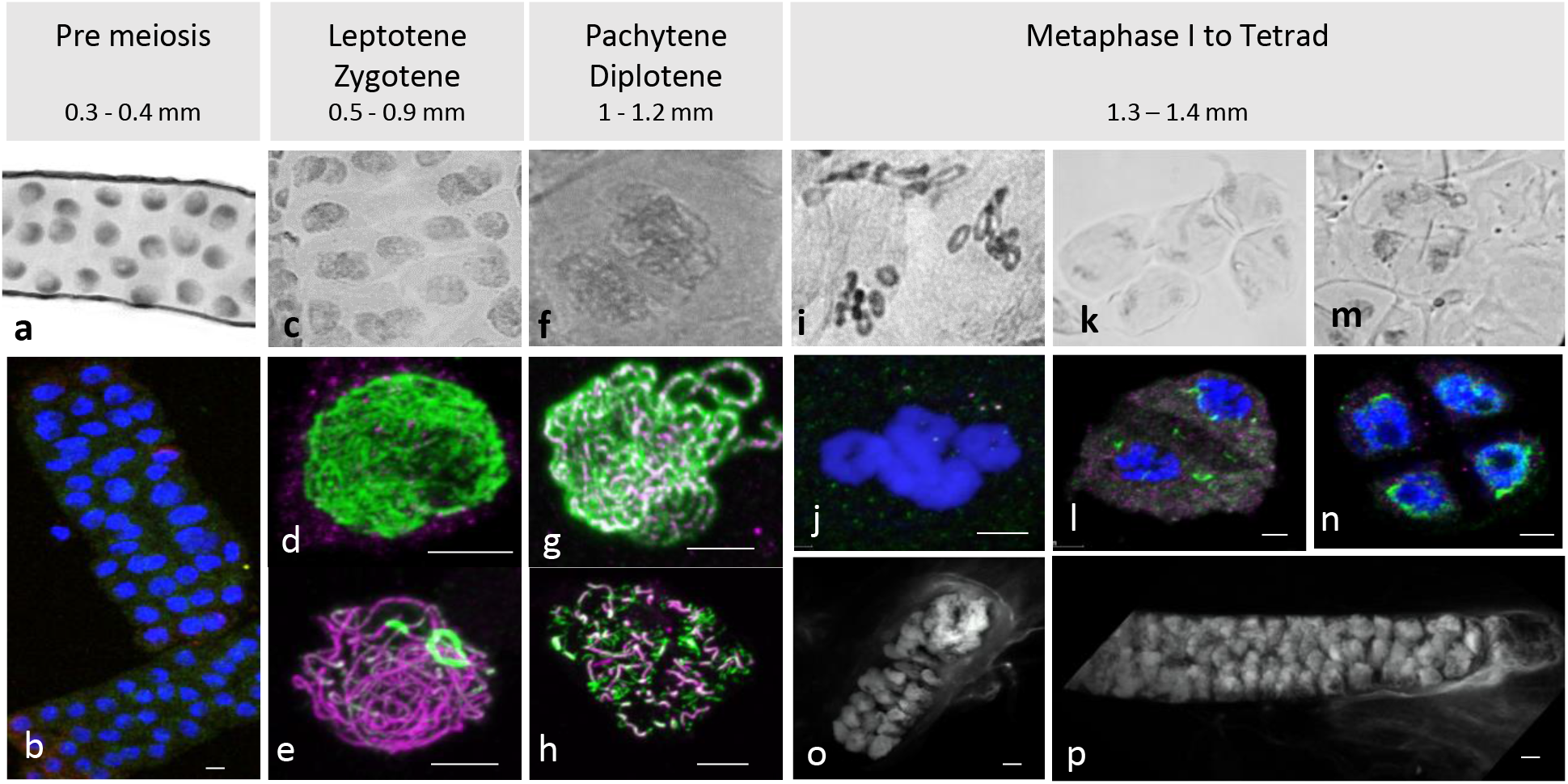
Anther and meiocyte collection and staging. Anthers were collected according to their size and staged with both aceto-carmine **(a,c,f,i,k,m)** and Immunocytology **(b,d,e,g,h,j,l,n). (a,b)** premeiotic stage; **(c)** leptotene/zygotene stage **(d);** leptotene stage; **(e)** zygotene stage; **(f)** pachytene stage; **(g)** pachytene stage; **(h)** diplotene stage; **(i,j)** metaphase I; **(k,l)** anaphase I; **(m,n)** tetrads. **(o, p)** Isolated fresh meiocytes bag at Leptotene/zygotene and pachytene/diplotene respectively. Aceto-carmine (Grey), DAPI (Blue and white for images **o** and **p**), ASY1 (Green), ZYP1 (Magenta). Scale bar 5μm Note images **(o)** and **(p)** have associated videos.

Acetocarmine stained pre-meiotic spreads showed small nuclei (Figure 1a) without any labelling by antibodies against the SC proteins HvZYP1 and TaASY1, although a signal can be detected with anti-TaASY1 mainly in the cytoplasm (Figure 1b). Contrary to the pre-meiotic stages, anther size and acetocarmine staining were not entirely reliable for leptotene–zygotene staging (Figure 1c) but immunostaining with antibodies against TaASY1 and HvZYP1 allowed us to easily distinguish between these two close developmental stages. At leptotene, linear ASY1 axes are formed while there is no or little labelling of HvZYP1 at the initiation of synapsis (Figure 1d). Zygotene on the other hand was determined by the presence of HvZYP1 labelling during synapsis progression as previously described (Colas *et al.*, 2017) (Figure 1e). Pachytene and diplotene were somewhat easier to find based on anther size and acetocarmine spreads (Figure 1f) and were distinguished using antibodies against TaASY1 and HvZYP1 proteins (Figure 1g–h). Pachytene (Figure 1g) was characterized by the polymerization of HvZYP1 between the ASY1 axes along the entire chromosomes and diplotene (Figure 1h) was identifiable by the appearance of tinsel chromosomes as previously described (Colas *et al.*, 2017). Since our primary focus was on synapsis and recombination, later stages from metaphase I to tetrads were combined into a single sample (Figure 1). Metaphase I is characterised by the presence of seven ring bivalents (Figure 1i, j) while anaphase I (Figure 1k, l) and anaphase II samples show chromosomes or chromatin segregation, respectively (Colas *et al.*, 2017). Anther samples of these later stages were not as synchronised as at prophase (leptotene–diplotene), containing both metaphase I and anaphase I or both anaphase II and telophase II. Very few anthers contained tetrads (Figure 1n) and none reached pollen stage.

A subset of anther samples that were staged as leptotene–zygotene and pachytene–diplotene by immuno-cytology were used to release meiocyte bags onto microscope slides. A sample of these were stained with DAPI to evaluate the quality of the preparation (Figure 1o–p; Supplemental Movies 1 and 2) and the remaining material transferred into Eppendorf tubes containing TRIzol® for subsequent total RNA isolation.

Altogether, the combination of anther size and immuno-cytology allowed us to delineate different developmental stages. We extracted total RNA and sequenced 18 samples (3 biological replicates of 2 meiocyte and 4 anther stages) generating a total of 2.1 billion reads from anther/meiocyte samples. For comparative purposes we extracted and sequenced RNA from 4 replicate germinating embryo (EMB) samples generating over 245 million reads.

### RNA-seq reveals abundant differential expression

For RNA-seq and differential gene expression analysis we first developed a reference **B**arley **An**ther **Tr**anscriptome (BAnTr). We did this for two important reasons. First, we anticipated that anther/meiocyte tissues potentially contain a set of very specific transcripts. So, to complement the Illumina short read RNA-seq data, we generated and included three 5’ cap- and 3’ polyA-captured PacBio IsoSeq datasets from a mixture of the RNA samples mentioned above. Second, we are working with the transformation reference barley cultivar Golden Promise and had recently generated a Golden Promise genome reference assembly (Schreiber *et al.*, 2020). Building a reference-guided transcriptome based on the same cultivar will be more complete and more accurate as cultivar specific transcripts will be included. To include as many barley transcripts as possible, the Illumina RNA-seq and IsoSeq reads of anthers and meiocytes were combined with the short reads from the germinated embryo samples and ultimately with the BaRTv1-Reference Transcript Dataset reported recently by Rapazote-Flores *et al.* (2019). The final transcriptome, BAnTr, comprised 65,795 genes and 119,778 transcripts.

For each sample, transcript quantification files were generated using Salmon (Patro *et al.*, 2017) in conjunction with BAnTr. The output quantification files were then read into the 3D RNA-seq pipeline (Guo *et al.*, 2019) to generate read counts which were converted to transcript per million (TPM) using the tximport R package (Soneson *et al*., 2016). The data used as input for the 3D RNA-seq pipeline has been deposited at figshare: https://doi.org/10.6084/m9.figshare.11974182. Read counts and TPMs were pre-processed and filtered to reduce noise and technical variance by excluding low abundance transcripts (keeping transcripts with a count per million reads ≥ 1 in at least 3 samples, and genes where at least one transcript passed the filtering). After filtering, 50,861 transcripts encoded by 31,918 genes remained. To analyse the relationship between replicates, a multidimensional scaling plot was generated using the transcript expression data showing clear separation between samples (Supplemental Figure 1). We set up the following comparisons to study differential expression: anthers at leptotene–zygotene *vs*. anthers at pre-meiosis; anthers at pachytene–diplotene *vs.* anthers at leptotene–zygotene; anthers at metaphase I–tetrad *vs.* anthers at pachytene–diplotene; meiocytes at leptotene–zygotene *vs.* anthers at leptotene–zygotene; meiocytes at pachytene–diplotene *vs.* anthers at pachytene–diplotene; and meiocytes at leptotene–zygotene *vs.* meiocytes at pachytene–diplotene. We defined differentially expressed genes (DEGs) as those showing a statistically significant (Benjamini-Hochberg corrected probability value ≤ 0.01) log_2_-fold change (log_2_FC) in expression ≥ 1 or ≤ −1 in stage and tissue comparisons. Using these criteria, a total of 10,713 DEGs (Supplemental Figure 2) were identified and for further analysis divided into three categories based on protein-coding potential: protein-coding, long non-coding RNA (lncRNA), and unclassified.

### Weighted gene co-expression analysis highlights gene co-expression modules relevant to meiotic processes

In addition to DEG analysis, we used the weighted gene co-expression network analysis (WGCNA) tool (Langfelder and Horvath, 2008) to identify co-expression networks within all 31,918 expressed genes in the anther and meiocyte dataset. The analysis identified 17 gene clusters (shown as colour-coded modules in Supplemental Figure 3) with an additional un-clustered grey module (Supplemental Table 1). The number of genes in each module ranged from 38 to 6209 (Supplemental Table 1). Variation was also evident in differential eigengene network analysis showing levels of correlation between clusters (Supplemental Figure 4). WGCNA navy and yellow modules were enriched in the meiocyte samples in comparison to anthers. In total, four modules (navy, yellow, green, and olive) showed higher expression in meiocytes in comparison to anthers (Figure 2). Gene ontology (GO) enrichment analysis highlighted mRNA and RNA binding GO terms as significantly enriched in the navy module (P value = 6E-16 and P value = 3.1E-13); the yellow module was significantly enriched for DNA binding (P value < 1E-30), RNA modification, and nucleosome GO terms (Figure 3) (P value < 1E-30 and P value < 1E-30). From the remaining modules, genes clustered in the red module showed a pattern of expression which was approximately inverse to genes clustered in the navy module (Supplemental Figure 3). The red module is significantly enriched in genes located in chloroplast components (Supplemental Figure 5, Supplemental Data set 1) (chloroplast thylakoid membrane, P value = 2.2E-13 and chloroplast envelope, P value = 3.7E-6). The blue and the light-green modules show a similar downward trend of expression with meiotic progression (Supplemental Figure 3).

**Figure 2:**
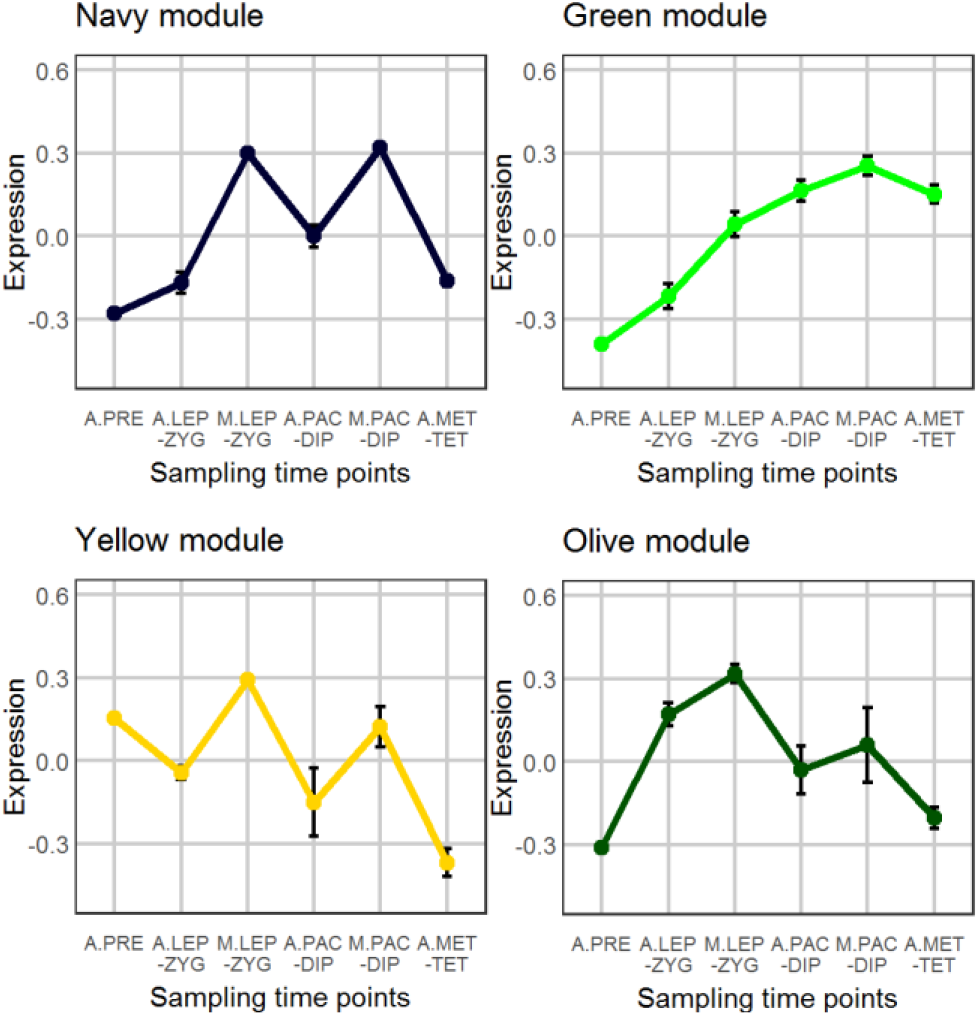
WGCNA analysis of co-expressed genes. A total of 17 modules were found and the selected four show an interesting pattern for meiocyte enriched genes.

**Figure 3:**
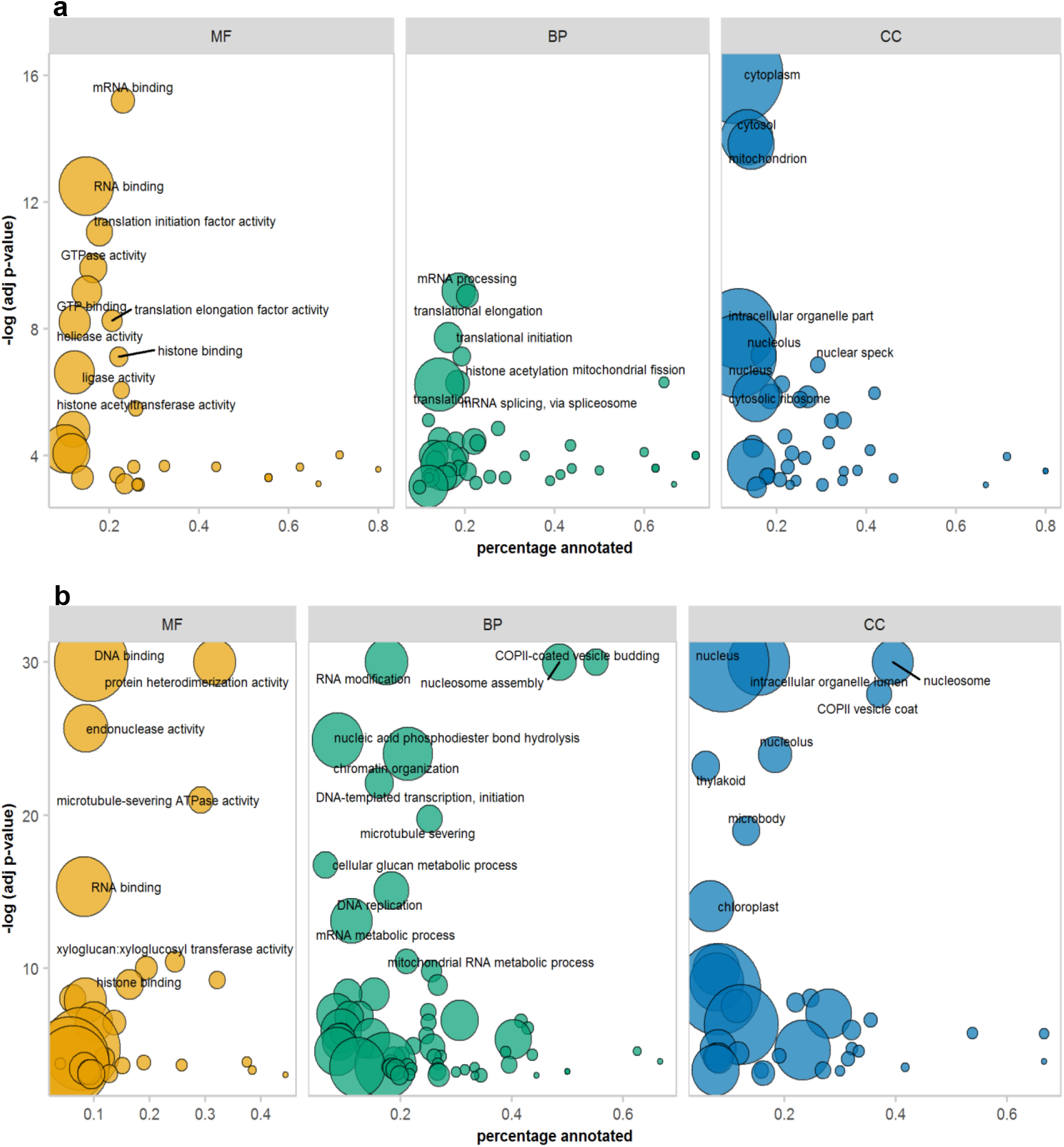
Gene ontology enrichment analysis. Results from the navy (a) and the yellow (b) modules of the WGCNA. Size of bubbles correspond to total number of proteins associated with the GO term. MF = molecular function, BP = biological process, CC = cellular component.

### Large scale transcriptional re-programming between pre-meiosis and leptotene–zygotene is largely stable throughout prophase I

3D RNA-seq analysis revealed a dramatic alteration in gene expression in early prophase I that largely stabilised as anther and meiotic tissues progress beyond leptotene–zygotene, with genes either continuing up-/down-regulation or showing no further significant changes (Figure 4a–c; Supplemental Table 2). The number of DEGs was highest (n=6119) in comparison of anthers at leptotene–zygotene versus pre-meiosis. Strikingly, 48% of the total number of DEGs and 65% of the down-regulated DEGs are lncRNAs in this comparison (Figure 4a, b). Compared with germinating embryos, anthers contain a much larger number of lncRNAs (Figure 5a). As expected, the majority of these lncRNAs are relatively short (Figure 5c) and comprise a single exon (Figure 5d). In anther tissues, there was a further pronounced, but much smaller, change in gene expression at pachytene–diplotene versus leptotene–zygotene (Figure 4a, b). However, gene expression in meiocytes appears to have already stabilised, with a meagre 4 DEGs (2 up-regulated; 2 down-regulated) between meiocytes at pachytene–diplotene versus leptotene–zygotene (Figure 4c). Only 164 genes (163 up-regulated; 1 down-regulated) are significantly differentially expressed in anthers at metaphase I–tetrad compared to pachytene–diplotene (Figure 4a, b). When lncRNAs are excluded from the analysis, the largest number of DEGs occurs in meiocytes at leptotene–zygotene when compared to anthers at the same stage (n=3517; Figure 4c); the number of protein-coding DEGs in comparison of anthers at leptotene–zygotene *vs*. pre-meiosis is 2710 (Figure 4b).

**Figure 4:**
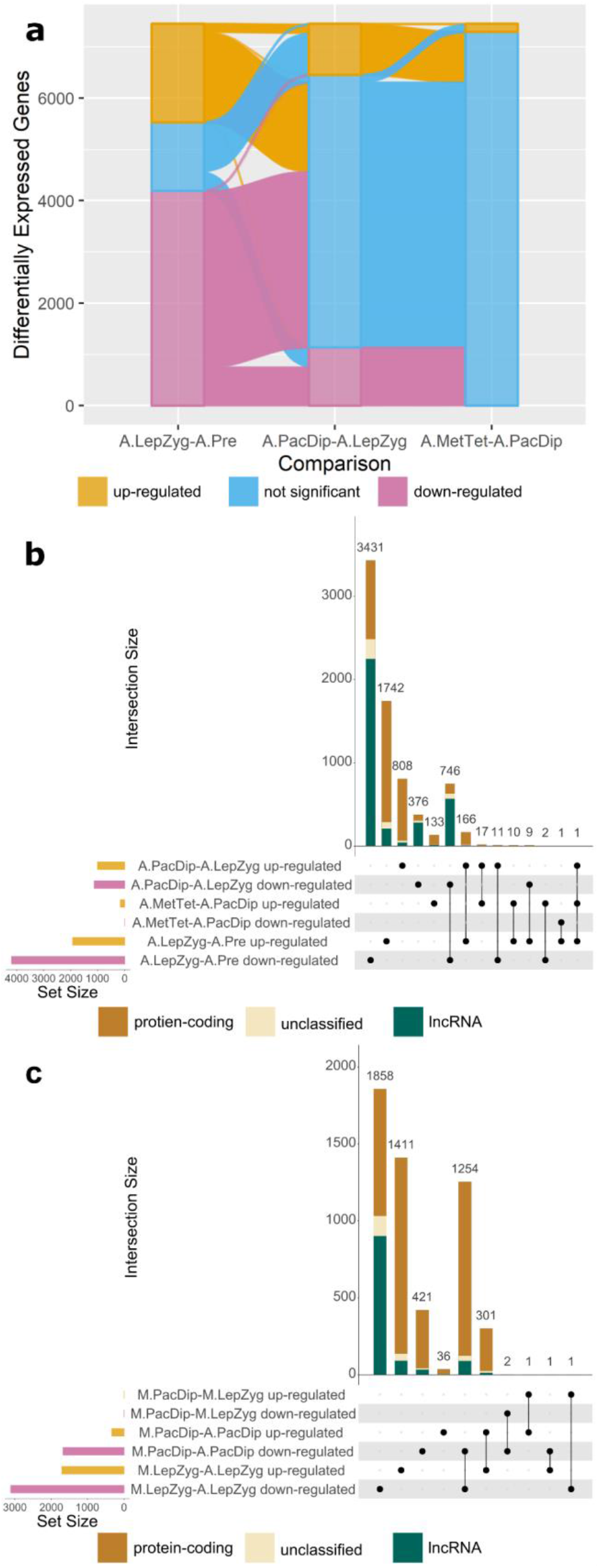
Comparisons of differential gene expression. **a)** Alluvial plot of significantly differentially expressed genes in anther (A) meiotic stage comparisons. The comparisons are: **A.LepZyg-A.Pre**, leptotene–zygotene versus pre-meiosis; **A.PacDip-A.LepZyg**, pachytene–diplotene versus leptotene–zygotene; and **A.MetTet-A.PacDip**, metaphase I–tetrad versus pachytene–diplotene. **b)** intersect plot of the same anther meiotic stage comparisons and **c)** intersect plot of meiocyte (M) stage and meiocyte-anther tissue comparisons. The combination matrix at the bottom of b and c plots indicates comparison-specific (left) and shared (right) subsets and the bar above it shows their sizes. The set size of each comparisons is shown on the left. The comparisons are: **M.PacDip-M.LepZyg**, meiocytes at pachytene–diplotene versus meiocytes at leptotene–zygotene; **M.LepZyg-A.LepZyg**, meiocytes at leptotene–zygotene versus anthers at the same stage; **M.PacDip-A.PacDip**, meiocytes at pachytene–diplotene versus anthers at the same stage.

**Figure 5:**
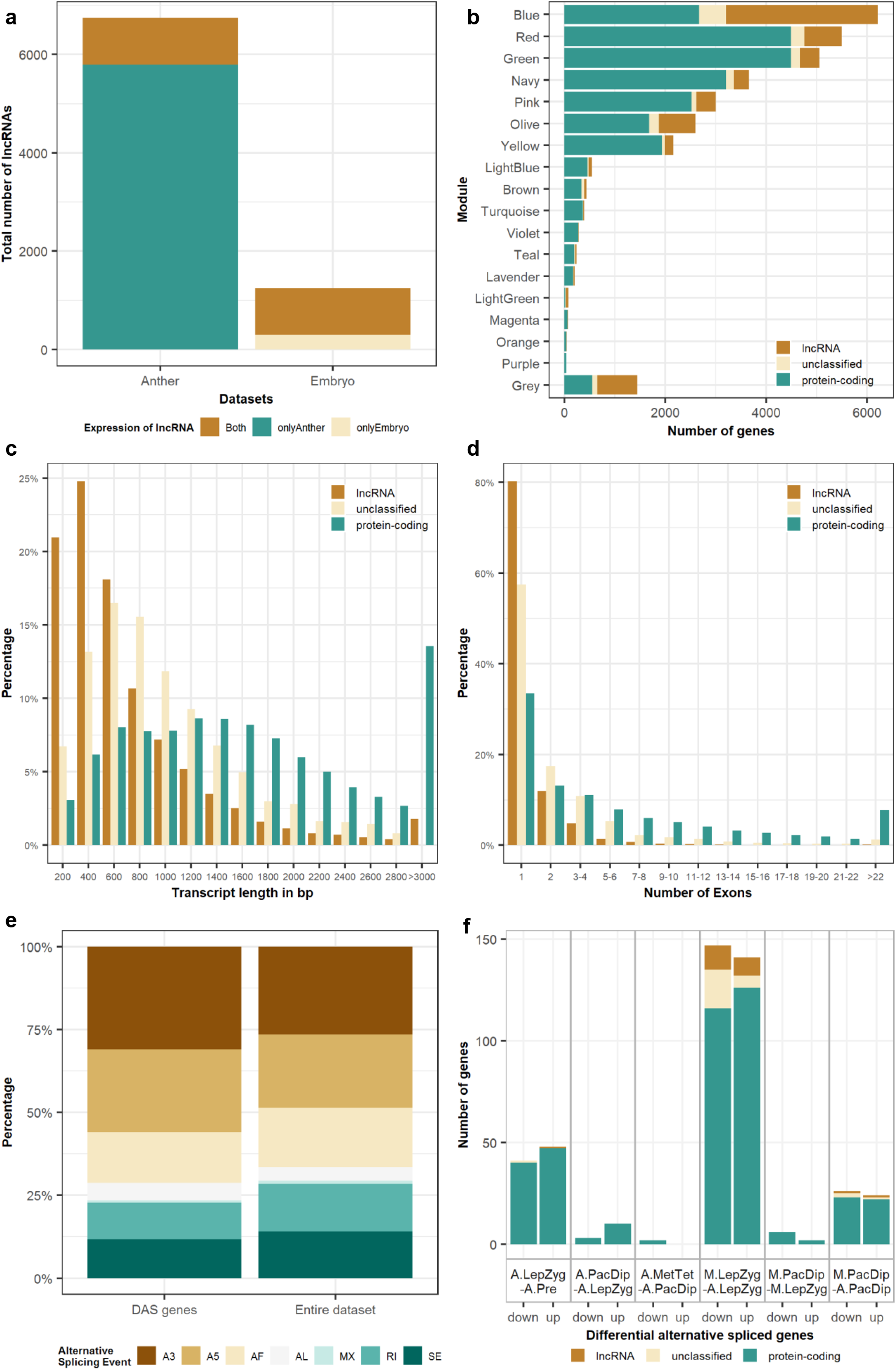
Differential expression and alternative splicing of different gene categories. **a)** Compared with germinating embryos, anthers are enriched in lncRNAs. **b)** Distribution of coding, noncoding and undefined genes in different modules. **c)** Length and **d)** exon number distributions of different transcript categories. e) Distribution of alternative splicing events for the entire dataset and the DAS (differential alternative spliced) genes. A3 – alternative 3’ splice-site; A5 – alternative 5’ splice-site; AF – alternative first exon; AL – alternative last exon; MX – mutually exclusive exons; RI – retained intron; SE – skipping exon. **f**) differential alternative spliced genes. The comparisons are: **A.LepZyg-A.Pre**, anther leptotene–zygotene versus anther pre-meiosis; **M.LepZyg-A.LepZyg**, meiocyte leptotene–zygotene versus anther leptotene–zygotene; **A.PacDip-A.LepZyg**, anther pachytene–diplotene versus anther leptotene–zygotene; **M.PacDip-A.PacDip**, meiocyte pachytene–diplotene versus anther pachytene–diplotene; **A.MetTet-A.PacDip**, anther metaphase I–tetrad versus anther pachytene–diplotene.

### Differential alternative splicing

A striking feature of many of the genes that are putatively involved in meiosis and recombination in barley is the number of introns and exons they contain. The average of our protein-coding transcripts in the BAnTr transcriptome is 7.5 exons per gene. Our selection of 121 potential meiotic genes has an average of 25 exons per gene—more than three times the average for all genes. One possible reason may be that regulation of these genes occurs at the post-transcriptional level through delayed pre-mRNA processing or differential alternative splicing (DAS). Indeed, 84 out of the 121 potential meiotic genes have more than one isoform. However, only two of those genes are significantly differentially alternatively spliced (HvHSP90.7 and HvPTB1a). The analysis of DAS genes across the whole transcriptome revealed that most contain isoforms with different 3’ or 5’ splice-sites, representing 48% of the total identified events (Figure 5e). Only 0.8% of the genes contained isoforms with mutually exclusive exons. The DAS genes were similarly distributed with no splicing event being strongly enriched in comparison to the whole dataset. GO enrichment analysis showed for both groups of DAS genes, i.e. with alternative 3’ or 5’ splice-sites, an enrichment in ribonucleoprotein complex (P value = 2.8E-4 and P value = 2.6E-5) which contains the small nuclear ribonucleoprotein particles (snRNPs) forming the spliceosome. We also observed enrichment of the GO terms U2-type prespliceosome (P value = 7.3E-5) and U1 snRNP (P value = 1.3E-4). A close look at the gene list confirmed both U2A-factor (BAnTr.GP.5HG019656) and U1-70k (BAnTr.GP.1HG004326, BAnTr.GP.1HG006488) components plus additional genes involved in forming the small nuclear ribonucleoprotein particle (snRNP). The most differentially alternatively spliced genes can be observed in the comparison of meiocytes to anthers in leptotene–zygotene stage (Figure 5f). GO enrichment analysis of down-regulated genes highlighted the same GO terms as above (U2-type prespliceosome, P value = 7E-5 and U1 snRNP, P value = 1.1E-4). The up-regulated genes were enriched in the GO term regulation of DNA damage checkpoint (P value = 2.5E-4).

### Genes annotated with roles in post-transcriptional and post-translational modification are a major component of up-regulated genes in prophase I

GO enrichment analysis of DEGs points to a sizeable role for RNA and post-translational modifications in prophase I (Supplemental Data set 2). This was most acutely highlighted in the comparison of meiocytes to anthers at leptotene–zygotene—representing the largest contrast in expression of protein-coding genes. Lys48-specific deubiquitinase activity (GO:1990380; P value = 0.00083), histone lysine demethylation (GO:0070076; P value = 0.00068), histone H3-K9 demethylation (GO:0033169; P value = 0.00074), ligase activity (GO:0016874; P value = 0.0002), and RNA modification (GO:0009451; P value = 3.60E-10) were all significantly enriched in this comparison. Enrichment of the RNA modification GO term reflected up-regulation of genes annotated as pentatricopeptide repeat (PPR) proteins, which regulate gene expression at the RNA level (Manna, 2015). Up-regulation of these genes also drives significant enrichment of endonuclease activity (GO:0004519; P value = 0.00088) in this comparison. Of the two up-regulated DEGs in meiocytes at pachytene–diplotene *vs.* leptotene–zygotene one was annotated as a PPR protein, the other was unannotated, and both down-regulated DEGs in this comparison were lncRNAs. Continued up-regulation of these genes at pachytene–diplotene in meiocytes *vs*. anthers at the same stage is also reflected in significant enrichment of mRNA binding (GO:0003729; P value = 0.00033). Up-regulated DEGs in this contrast group were enriched in helicase activity (GO:0004386; P value = 0.00088) and synapsis (GO:0007129; P value = 0.00023), reflecting the formation of the synaptonemal complex in meiocytes at this stage. GO enrichment of ligase activity in up-regulated DEGs in meiocytes at leptotene–zygotene compared to anthers at the same stage is driven by up-regulation of E3 ubiquitin ligases. In total, 890 ubiquitin or ubiquitin-like (SUMO, NEDD8) E3 ligases were annotated in the BAnTr transcriptomic dataset; 133 of these were specific to anther and meiocyte tissues, 63 were unique to germinating embryos, and 589 were expressed in both anther and germinating embryo tissues. 166 of these were differentially expressed in a least one contrast group; 71 were up-regulated in meiocytes compared to anthers at the same stage (Supplemental data set 3). Notably, those up-regulated in meiocytes included many E3 ligases—or multi-subunit E3 ligase components—with confirmed roles in meiosis in other organisms. These included: a SKP1 orthologue (BAnTr.GP.5HG012508) and three F-box proteins (BAnTr.GP.2HG002834, BAnTr.GP.3HG014344, BAnTr.GP.3HG014348) which form part of the multi-subunit SKP Cullin F-box (SCF) E3 ligase complex; one RING-H2 component of the anaphase promoting complex (BAnTr.GP.1HG006974); and three seven-in-absentia (SINA) E3 ligases (BAnTr.GP.2HG018292, BAnTr.GP.3HG000774, BAnTr.GP.3HG000850). HEI10, a highly conserved E3 ubiquitin ligase which is known to be involved in DSB repair (Ziolkowski *et al.*, 2017), was stably expressed in all tissues and stages, including in germinating embryo tissues. Although no NEDD8 E3 ligases are differentially expressed in any stage or tissue comparison, NEDD8-specific protease activity (GO:0019784; P value = 0.00029) is enriched in pachytene–diplotene compared to leptotene–zygotene in anthers. Significant up-regulation of Lys48 specific deubiquitinase activity, in parallel to E3 ligase activity, might reflect delicate regulation of the ubiquitination cascade at this stage. However, of the 4 significantly up-regulated protein-coding genes assigned this GO by PANNZER2, only one (BAnTr.GP.7HG015226) is explicitly annotated as a deubiquitinating enzyme (DUB); two are annotated as B-box transcription factors and the other as an O-fucosyltransferase. Lys-48 deubiquitination (GO:0071108; P value = 3.20E-5) is also enriched in pachytene–diplotene compared to leptotene–zygotene in anthers reflecting the up-regulation of four protein-coding genes annotated as otubain-like DUBs, active on Lys48-linked polyubiquitin chains and NEDD8 (Edelmann *et al.*, 2009).

Histone lysine demethylation and histone H3-K9 demethylation enrichment in meiocytes compared to anthers at leptotene–zygotene is driven by up-regulated protein-coding genes orthologous to argonaute (AGO) proteins and demethylases KDM3 (BAnTr.GP.1HG003366), PKDM9 (BAnTr.GP.UnG004036), JMJ705 (BAnTr.GP.UnG004032 & BAnTr.GP.UnG004334), and JMJ706 (BAnTr.GP.1HG007084). Indeed, of 21 AGO orthologues, 10 are differentially expressed in at least one stage or tissue comparison. When the log_2_FC cut-offs for differential expression are disregarded 14 AGO orthologues are significant in at least one contrast group.

Protein-coding genes which are up-regulated in anthers *vs*. anthers at successive stages and down-regulated in meiocyte *vs*. anther contrast groups of the same stage are largely reflective of water transport, photosynthesis, cell signalling, amino acid transport, and other metabolic processes (Supplemental Data set 3). However, down-regulated DEGs in anthers at leptotene–zygotene vs. pre-meiosis are more informative. The switch to prophase I in anthers, in addition to the large-scale down-regulation of lncRNAs, coincides with massive down-regulation of protein-coding genes annotated with RNA-directed DNA polymerase activity (GO:0003964; P value < 1E-30) all of which are expressed only in anther tissues and not in germinating embryos. These also formed a sizeable component of the down-regulated DEGs annotated as nucleic acid binding (GO:0003676; P value < 1E-30) and zinc binding (GO:0008270; P value < 1E-30) proteins; 89% of both groups were also anther specific. DNA binding transcription factors (GO:0003700; P value = 3.20E-5) and transcriptional regulation related GO terms GO:0017053 and GO:0006355 (P values = 5.50E-5 and 2.20E-6, respectively) were also enriched in down-regulated DEGs in this contrast group.

### Not all meiotic genes are expressed as expected—but some are

We probed pre-meiotic and meiotic cells with antibodies against two synaptonemal complex (SC) (ASY1 and ZYP1) proteins and one recombination (DMC1) protein. The protein products of *HvASY1* and *HvZYP1* were detected during prophase I but not in pre-meiotic cells (Figure 6A). HvDMC1 detection on the other hand shows a diffuse, mostly cytoplasmic, signal in pre-meiotic cells. During meiosis, HvDMC1 appears as discrete foci in leptotene–diplotene nuclei before becoming diffuse again at metaphase I–tetrad. All three genes are transcribed at pre-meiosis (Figure 6B) and belong to the navy WGCNA module showing higher transcript abundance in prophase I meiocytes (Figure 2). We then analysed the gene-level expression of 121 known meiotic genes which were distributed among 11 different modules. While all were expressed, only 29 were differentially expressed across the samples using our experimental thresholds (Figure 6B, Supplemental Figure 6). It is noticeable that the overall level of expression of some genes, like *HvPCNA* and members of the *HSP90* family, remains high in all stages and tissues while others, like *HvMET1b*, *HvMRE11B*, *HvXRCC3* and *HvDDM1C*, remained relatively low.

**Figure 6:**
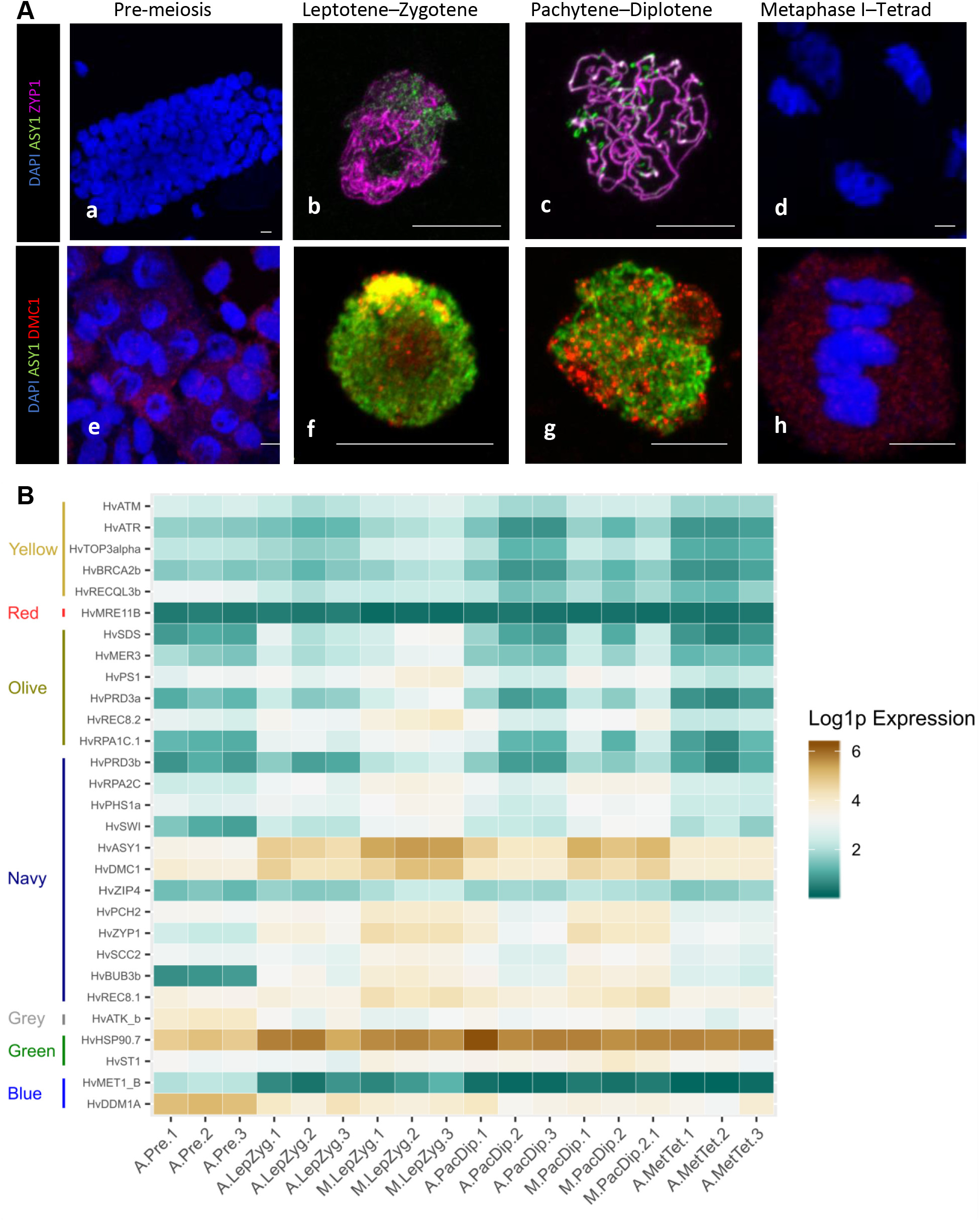
Expression of selected meiotic genes. **A)** immuno-staining of meiotic nuclei at four developmental stages for HvZYP1 (magenta, a-d), and HvDMC1 (red, 2-h) proteins. All samples were stained with anti-ASY1 antibody (green) and counterstained with DAPI (Blue). Scale bar 10 mm. **B)** Heatmap expression profile of meiotic genes with a statistically significant log fold change in at least one tissue or stage comparison. The genes are ordered and grouped by WGCNA module on the vertical axis. Genes were extracted from the total dataset, transcript counts log transformed, and plotted using ggplot2 (Wickham, 2016) in R (script available at https://github.com/BioJNO/BAnTr). The samples (3 replicates each) are A.PRE, anther pre-meiosis; A.LEP-ZYG, anther leptotene–zygotene; A.PAC-DIP, anther pachytene–diplotene; A.MET-TET, anther metaphase I–tetrad; M.LEP-ZYG, meiocyte leptotene–zygotene; M.PAC-DIP, meiocyte pachytene–diplotene.

Of the 29 meiotic genes, *HvASY1*, *HvDMC1, HvZYP1*, *HvBUB3b*, *HvSDS*, *HvSWI*, *HvRPA2c*, *HvRPA1c.1*, and *HvHSP90.7* were significantly up-regulated from pre-meiosis to leptotene–zygotene. Further gene-level expression comparisons showed 19 putative meiotic genes (*HvASY1*, *HvATM*, *HvATR*, *HvBRCA2b*, *HvBUB3b*, *HvMER3*, *HvPCH2*, *HvPHS1a*, *HvPRD3a*, *HvPRD3b*, *HvPS1*, *HvREC8.1*, *HvREC8.2*, *HvRPA2C*, *HvSCC2*, *HvSWI*, *HvTOP3alpha*, *HvZYP1*, and *HvZIP4*) significantly up-regulated in meiotic cells compared to anthers at the same stage. In comparison to our control dataset of germinating embryo samples, expression of only 6 of these meiotic genes are specific to anthers. However, while the rest are expressed in the EMB samples, 16 show lower gene expression in this tissue. The other nine genes show EMB gene expression similar to at least one anther developmental stage.

### Transcriptional regulation

Anther and meiocyte transcriptomes have been determined in several plant species but the mechanisms of gene regulation in these tissues are still poorly understood. The varied gene-level expression patterns of modules generated by WGCNA analysis in this dataset could be explained by steady-state gene expression regulation, including at the transcriptional level. Using the Plant Transcription Factor (TF) Database (Pérez-Rodríguez *et al.*, 2010), we found a total of 1353 annotated TFs of which 382 (28.2%)—belonging to several different TF families (Supplemental Dataset 4)—were differentially expressed in at least one stage or tissue comparison. 140 of these were expressed in anther tissues and not in germinating embryos. The largest families in the significant log_2_FC subset are bHLH (n=29) and MYB (n=29), (Figure 7a). The distribution of TFs families in different expression modules was determined with the red module containing the highest number (n=154; Supplemental Dataset 5), a module showing down-regulation in meiocytes when compared to anthers at both stages of leptotene–zygotene and pachytene–diplotene (Supplemental Figure 3). This is reflected in the significant GO enrichment of transcriptional regulation in anthers at leptotene–zygotene *vs.* pre-meiosis. 254 (66.5% of transcription factors significant in any contrast group) show differential expression in comparisons between meiocytes and anthers at either pachytene or leptotene, with 170 and 84 TFs significantly different in one or both stages, respectively (Figure 7b).

**Figure 7:**
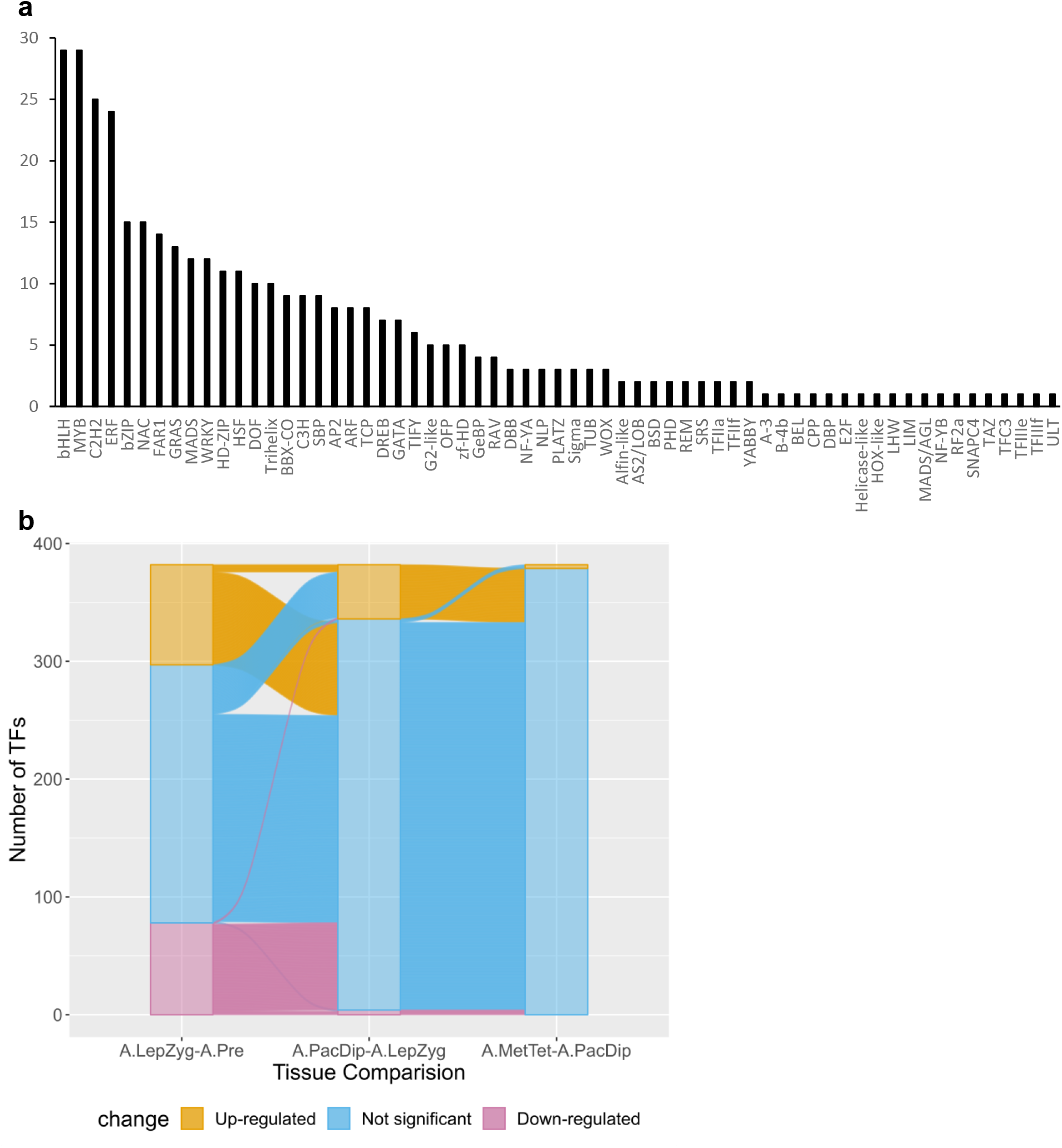
Expression of transcription factor (TF) families in anthers and meiocytes. **a)** Total number of TFs per family that are expressed in anthers and meiocytes; **b)** Number of differentially expressed TFs determined by comparing their transcript levels in anthers at different stages. The comparisons are: **A.LepZyg-A.Pre**, anther leptotene–zygotene versus anther pre-meiosis; **A.PacDip-A.LepZyg**, anther pachytene–diplotene versus leptotene–zygotene; versus meiocyte leptotene–zygotene; **PAC vs A.LEP**, anther pachytene–diplotene versus anther leptotene–zygotene; **A.PAC vs M.PAC**, anther pachytene–diplotene versus meiocyte pachytene–diplotene; **A.MetTet-A.PacDip**, anther metaphase I–tetrad versus anther pachytene–diplotene.

## DISCUSSION

Despite technical advances, and the importance of the reproductive phase in crop breeding, the regulation of the meiotic transcriptome in plants is still poorly understood (Zhou and Pawlowski, 2014). Using microarray and RNA-seq platforms, several previous studies have highlighted the developmental changes of the anther transcriptome in several species (Dukowic-Schulze and Chen, 2014). However, most of these studies were limited to early meiosis (Dukowic-Schulze *et al.*, 2014) or even focused on mitotic–meiotic transition stage only (Yuan *et al.*, 2018; Nelms and Walbot, 2019). In this study, we sought to establish the most comprehensive transcriptome of barley anthers and meiocytes spanning pre-meiosis–tetrad stages. The barley cultivar Golden Promise (GP) was chosen as it is the reference cultivar for barley transformation which is relevant for future functional studies. In addition, we recently sequenced its genome (Schreiber *et al.*, 2020), which will assist with interpretation of the transcriptome data, and developed a TILLING population using ethyl methanesulfonate (Schreiber *et al.*, 2019). We used immuno-cytology to carefully stage all anther and meiocyte samples and analysed a combination of Illumina and PacBio reads using the most up to date bioinformatic pipeline. This allowed us to identify novel transcripts and study transcriptome dynamics throughout meiosis.

### lncRNAs play a significant role at prophase I entry

The number DEGs in staged anther tissues declines from its highest level between leptotene–zygotene and pre-meiosis with the largest down-regulation of genes of any comparison. This supports previous studies in *Arabidopsis* (Chen *et al.*, 2010; Yang *et al.*, 2011), maize (Dukowic-Schulze *et al.*, 2014) and sunflower (Flórez-Zapata *et al.*, 2014; Flórez-Zapata *et al.*, 2016) showing large scale transcriptional re-organisation at meiosis entry. A strikingly large component of differentially expressed genes at this transition were lncRNAs—98% of which were expressed in anthers but not in germinating embryos. High expression of lncRNAs in plant reproductive tissues has been described in both sunflower (Flórez-Zapata *et al.*, 2016) and in maize (Li *et al.*, 2014). Flórez-Zapata *et al.* (2016) reported large scale differential expression of lncRNAs in sunflower meiocytes when contrasted with somatic tissues; and between meiocytes in comparison of genotypes exhibiting significantly different recombination rates. Our results indicate that, similar to sunflower, meiocyte specific lncRNAs may indeed play an important role in prophase I in barley. However, our results also indicate that expression of many of these lncRNAs is down-regulated during the transition to prophase I—indicating that their most important functional roles may be prior to meiosis. LncRNAs are emerging as key plant development regulators (Ariel *et al.*, 2015; Shafiq *et al.*, 2016) and their role in animal and plant sexual reproduction is now well established (Golicz *et al.*, 2018). In yeast, they promote pairing of homologous chromosomes (Ding *et al.*, 2012). LncRNAs are increasingly considered precursors or target mimics of small RNAs (sRNAs), which are also starting to emerge as key players in gene regulation during meiosis. Different classes of phased small-interfering RNAs (phasiRNAs), for example, have been shown to be enriched in maize anthers at different meiotic stages (Zhai *et al.*, 2015) and shown to be present in most angiosperms (Xia *et al.*, 2019). Deep sequencing technologies have enabled global analysis of meiotic sRNA in sunflower showing sequence similarity to 40% of lncRNAs (Flórez-Zapata *et al.*, 2016). In *Arabidopsis*, Huang *et al.* (2019) revealed different meiotic and mitotic sRNA landscapes and found that meiocyte-specific sRNAs (ms-sRNAs) are significantly enriched in genic regions contrary to somatic small interfering RNAs that are enriched in intergenic regions. A high proportion of these ms-sRNAs (69%) were found to be DSB-dependent (Huang *et al.*, 2019).

### Significant post-transcriptional and post-translational regulation during prophase I

Comparison of meiocytes and anthers at leptotene–zygotene presented the greatest contrast in protein-coding gene expression. GO enrichment analysis of up-regulated DEGs in this contrast group suggests heightened importance of ubiquitination, RNA modification, and histone demethylation in prophase I regulation.

#### Post-transcriptional RNA modification through PPR proteins

Significant enrichment of the RNA modification GO term in meiocytes compared to anthers at leptotene–zygotene reflected several up-regulated DEGs annotated as PPR proteins. PPR proteins are found in all eukaryotes but are remarkably abundant in land plants, indicative of massive expansion during land plant evolution (Barkan and Small, 2014; Gutmann *et al.*, 2020). Through sequence-specific binding, PPR proteins can mediate RNA folding, splicing, degradation, cleavage, and editing (Barkan and Small, 2014). PPR proteins are associated with organelles, deleterious mutations in which lead to defects in photosynthesis or oxidative phosphorylation (Barkan and Small, 2014). Dukowic-Schulze *et al.* (2014) reported up-regulation of carbohydrate metabolism and mitochondrial genes in maize meiocytes compared to whole anther tissues and seedlings, arguing that this indicated high energy demand concomitant with chromosome movement in early prophase I. Up-regulation of PPR proteins in early barley prophase I may support this, indicating a general increase in mitochondrial activity in step with a detectable spike in PPR protein mediated RNA regulation.

#### Ubiquitin ligases and deubiquitinating enzymes

E3 ubiquitin ligases are an important regulatory component during meiosis in many organisms (Okamato *et al.*, 2012; Mohammad *et al.*, 2018). E3 ligases interact with target proteins to facilitate, directly or indirectly, modification of the substrate with ubiquitin—conferring substrate specificity to the ubiquitination cascade (Iconomou and Saunders, 2016). Several E3 ligases, and multi-subunit E3 ligase components, have been described with significant roles in meiosis in *Arabidopsis* (Yang *et al.*, 1999, Wang and Yang, 2006; Yang *et al.*, 2006), rice (He *et al.*, 2016; Zhang *et al.*, 2017), and wheat (Li *et al.*, 2006; Hong *et al.*, 2013). Further, enrichment of E3 ligases in pre-meiotic pollen mother cells has been reported in both rice (Tang *et al.*, 2010) and maize (Yuan *et al.*, 2018). Our findings suggest that ubiquitination through E3 ligase activity is of continued importance throughout prophase I. E3 ligase genes up-regulated in meiocytes include an ASK1 orthologue, an *Arabidopsis* S-phase kinase-associated protein 1 (SKP1) which interacts with cullin and F-box proteins to form the SKP-Cullin-F-box (SCF) E3 ligase complex (Yang *et al.*, 1999). SKP1-like proteins are functionally conserved in plants, with the wheat ASK1 equivalent, TSK1, able to partially rescue fertility in *Arabidopsis ask1* mutants (Li *et al*., 2006). ASK1 was identified as a negative regulator of male recombination, essential for the proper release of chromatin from the nuclear membrane (Wang and Yang, 2006; Yang *et al.*, 2006). Several F-box proteins, putatively interacting in the SCF E3 ubiquitin ligase complex, also displayed differential expression throughout prophase I. In rice, mutations in two F-box proteins interacting with the ASK1 equivalent OSK1, MOF and ZYGO1, have also been shown to result in male sterility (He *et al.*, 2016; Zhang *et al.*, 2017). Several genes annotated as SINA E3 ligases also displayed differential expression during prophase I. A recent study in *Drosophila melanogaster* females identified a SINA E3 ligase which regulated both assembly and disassembly of the SC, preventing aberrant polymerization and polycomplex formation of SC components (Hughes *et al.*, 2019). Although no such function of SINA E3 ligase is so far reported in plants, the E3 ligases highlighted in this work are attractive candidates to investigate a similar role for these proteins in regulating SC formation in barley. DUBs, particularly those targeting lys48 ubiquitin chains, are also differentially expressed throughout prophase I. DUBs facilitate removal of ubiquitin from proteins; in this way they reverse ubiquitination mediated by E3 ligases. Lys48 is one of seven lysine residues through which ubiquitin can form covalent C-terminally linked chains (Kulathu and Komander, 2012; López-Mosqueda and Dikic, 2014). Although targeting for proteasomal degradation is the canonical function of ubiquitination, first described by Ciehanover *et al*. (1978), many other effects of this modification have been observed such as recruitment of binding partners (Huang and D’Andrea, 2010), activation (Xu *et al*., 2009), or nuclear uptake (Plafker *et al*., 2004). The diversity of substrate fates is derived from diversity in chain topology (Komander and Rape, 2012). Simultaneous up-regulation of E3 ubiquitin ligase and DUB expression may represent both very tight control of substrate modification, where ligase and DUB build and remove the same chain topology respectively, and specific reduction in lys48 ubiquitination while alternative chain linkage types are enriched, where they do not.

#### Methylation, demethylation, and AGO genes

We checked the expression dynamics of sRNA biogenesis and AGO genes, for which protein products were detected in our recent anther proteomic study (Lewandowska *et al.*, 2019), and found the expression of Dicer-like 3 (DCL3) and 10 out of 21 AGO genes changed significantly during barley anther and meiocyte development (Supplemental Figure 7). This gene expression profile highlights the importance of sRNA biogenesis and their guiding AGO proteins during meiosis. However, further analysis of these pathways and their target loci is needed to fully describe their role in meiotic development. One such function has been demonstrated in a recent study by Walker *et al.* (2018) showing that *de novo* RNA-directed DNA methylation (RdDM) induces cell-lineage-specific epigenetic marks regulating meiotic gene expression in *Arabidopsis*.

mRNA methylation is emerging as another important level of meiotic gene regulation. The N6-methyladenosine (m^6^A) modification was shown to be associated with mRNA translatability in yeast (Bodi *et al.*, 2015) and *Xenopus* (Qi *et al.*, 2016). More recently, Bushkin *et al.* (2019) showed that m^6^A methylation at the 3’ UTR of the mRNA encoding Rme1p, a transcriptional repressor of meiosis, results in its degradation allowing meiosis initiation in yeast. Current epitranscriptome methods necessitate a large amount of mRNA making it difficult to implement in plant meiocytes. Instead, we determined transcript levels of known plant m^6^A pathway genes (Yue *et al.*, 2019), for which protein products were detected in our anther proteomic study (Lewandowska *et al.*, 2019). The expression of these genes was relatively low and stable throughout anther and meiocyte development. However, their translation status and potential function in these tissues remains to be elucidated. It will be interesting to compare the expression profiles of these enzymes in different plant species. Indeed, Liu *et al.* (2020) profiled m^6^A and N^6^,2’-O-dimethyladenosine (m^6^Am) across human and mouse tissues and suggested that the difference in these epitranscriptomic marks is greater between species than tissue types.

### Expression of meiotic genes does not always reflect the timing of their functional roles in meiocytes

Using phylogenomic analysis, we compiled a large inventory of barley orthologs of known meiotic genes. We determined their relative expression levels and found them to be distributed in 11 different expression modules. In total, 24% of selected meiotic genes were differentially expressed during anther and meiocyte development. Like previous studies, our transcriptomic data provided evidence that many known meiotic genes are transcribed prior to prophase I. However, the timing of their translation is still not fully resolved due to limited availability of antibodies (this work), issues related to sample staging, and proteomic resolution (Zhang *et al.*, 2014; Yuan *et al.*, 2018). Genome-wide ribosome profiling could be achieved by using transgenic tools like RiboTag/RNA-seq (Lesiak *et al.*, 2015). Such analysis was performed in mice revealing alteration of maternal mRNA translation in oocytes at meiotic re-entry (Luong *et al.*, 2020).

### Expression of transcription factors changes throughout anther and meiocyte stages

To decipher the mechanisms of meiotic transcriptome regulation, we analysed our transcriptomic data for transcriptional regulation. The involvement of many transcription factors (TFs) in transcriptome reprogramming during pre-meiotic anther differentiation has been shown in maize (Zhang *et al.*, 2014). This study was recently expanded by analysing TFs expression at tissue level using microdissection (Yuan *et al.*, 2018). However, both studies used microarrays, a closed technology, that may underestimate the number of TFs. We found a total of 1353 annotated TFs (63 families) of which 28.2% were differentially expressed in at least one stage or tissue comparison. In agreement with previous maize studies, our data also reveal major TFs expression changes occurring at and beyond mitotic–meiotic transition. Beside these global analyses, functional studies of TFs are still lacking to fully understand meiotic gene regulation. More recently, *TBP-ASSOCIATED FACTOR 4b* (*TAF4b*), encoding a subunit of the RNA polymerase II general transcription factor TFIID, was shown to be enriched in meiocytes and to control transcription of genes involved in the meiotic cell cycle and recombination in *Arabidopsis* (Lawrence *et al.*, 2019). In our study, a single barley TAF4b homolog gene (BAnTr.GP.6HG009004) was found in the green module showing increasing expression (P value <0.01) throughout anther and meiocyte stages, though below our threshold of log_2_FC >1 in all comparisons.

### Alternative splicing

Although alternative splicing (AS) is known to increase the genome coding capacity, its activity during meiosis has been studied as far as we are aware only in mouse testis (Schmid *et al.*, 2013) and yeast (Kuang *et al.*, 2017). Intron retention events were found to be enriched in both mouse (Naro *et al.*, 2017) and yeast meiocytes (Kuang *et al.*, 2017). They were also enriched during *Arabidopsis* flower development (Wang *et al.*, 2014). Here, GO enrichment analysis showed enrichment of terms that are connected to alternative splicing. This contained genes involved in forming the spliceosome or spliceosome-associated non-snRNP proteins. Contrary to our expectations we only identified two meiotic genes as significantly differentially alternatively spliced. In future we would like to extend our potential meiotic gene list as the 121 genes are only a small subset and need to be extended to potential orthologous genes identified in studies from mice, yeast, and *Arabidopsis*. We also might be able to extend the list and add barley specific meiotic genes through this dataset.

### Limitations and future work

Our analysis of barley anther and meiocyte transcriptomes provides evidence of a multi-faceted regulatory network orchestrating meiotic gene regulation. Sequencing of sRNA and DNA methylation in anther and meiocyte at pre-meiosis to leptotene–zygotene stages with the strongest transcriptomic changes will enhance the full picture of these interconnecting gene regulation mechanisms. Newly developed technologies, like single-cell RNA-seq (Denyer *et al.*, 2019) and spatial transcriptomics (Giacomello *et al.*, 2017), will increase the resolution of anther and meiocyte transcriptomes even at sub-cellular level. Nascent transcript sequencing by NET-seq (Churchman and Weissman, 2011), GRO-seq (Hetzel *et al.*, 2016) or Neu-seq (Szabo *et al.*, 2020) and ribosome profiling by RiboTag-seq (Lesiak *et al.*, 2015), for example, could be used to directly evaluate transcriptional and translational activities throughout anther and meiocyte development. The combination of immuno-cytological staging and RNA-seq data has allowed us to build a robust time-resolved barley anther and meiocyte transcriptomic dataset. We detected large scale down-regulation of lncRNAs at meiosis entry and enrichment of differential alternative splicing at early prophase I. In addition to changes in expression, we also revealed the diversity of transcription factors accompanied by several other post-transcriptional and post-translational regulatory networks. Our data will be informative for research in anther and meiocyte development in other plant species. In barley, our data has already contributed to building the first barley reference transcript dataset (BaRTv1). It will also be used in further proteomic studies of staged anthers and help in the annotation of barley genome and pangenome sequencing efforts.

## MATERIALS AND METHODS

### Plant material

Barley cv. Golden Promise plants were grown in cereal compost in a controlled growth room at 18°C for 16h light and 14°C for 8h dark and 70% humidity. When plants reached the desired stage (pre-meiosis/meiosis) at 5–7 weeks post-germination, they were processed individually by collecting their spikes at the last minute before anther sampling to minimise stress effect. Surfaces and tools were cleaned with 70% Ethanol and RNase AWAY® prior to material collection and all steps were done wearing gloves.

The collected spikes were placed in a Petri dish with wet filter paper (using RNase-free milli-Q water) on a cool pack or ice during the sampling, including under the stereomicroscope to avoid RNA degradation. Fine tweezers were used to remove awns and collect anthers of all florets between the fifth and fifteenth position from the bottom under a stereomicroscope resulting in 60 anthers on average. All materials were collected in the afternoon between 1:00 pm and 3:00 pm only to improve sample homogeneity. Four stages were collected: pre-meiosis/G2, leptotene/zygotene, pachytene/diplotene and all stages from metaphase I to tetrad.

Each barley floret contains 3 synchronized anthers. One anther was staged using acetocarmine staining and the two others were collected in 1.5 ml Eppendorf tube containing 100 μl RNA later. Anthers of the same length and meiotic stage were pooled together. From this tube, 10 random anthers were transferred into a new tube with 1x PBS buffer for staging validation with immunostaining using meiosis specific antibodies. The process was repeated until at least 3 replicates of 60 – 200 anthers were collected for each meiotic stage.

To isolate meiocytes, 30 to 50 anthers of the same length and meiotic stage based on acetocarmine staining were collected in 1.5 ml Eppendorf tube containing 0.01 M citrate buffer pH 4.5 and kept on ice for meiocytes isolation. Ten random anthers were transferred into a new tube with 1x PBS for further staging with specific antibodies. A maximum of 5 anthers were transferred onto a cavity slide with cold 5 – 10 μl of citrate buffer. The slide was placed on a cooling pack under the stereomicroscope for dissection. After removing their tips with insulin needles, anthers were tapped gently to release the meiocytes bag. Using RNase-free 10 μl tips, released meiocyte bags were transferred into a new 0.5 ml Eppendorf tube containing 10 μl of citrate buffer (with added RNAse Inhibitor) and anther tissue was discarded. When all the anthers were processed, an aliquot of 10 μl was transferred onto a slide and the sample was mounted in Vectashield® containing DAPI (H-2000, Vectorlabs) to check the quality of the meiocyte bags (Supplemental videos 1 and 2). The remaining meiocytes were topped with 50 μl TRIzol® and stored in the fridge until extraction.

For comparison we also included four samples of germinating embryo (EMB) as those have been shown to represent a wide range of genes (IBGSC, 2012). Barley cultivar (cv.) Golden Promise seeds were germinated in a Petri dish on wet paper for 5 days in dark. Three EMB were collected per sample by removing the residual seed material. In total four samples were collected and used for total RNA extraction.

### Cytology and Imaging

Ten anthers of each pool were fixed in 4% formaldehyde (1x PBS/0.5% TritonP™ X-100/0.5% Tween 20) for 20 minutes, rinsed twice in 1x PBS and ground gently in the tube with a disposable pestle. Tubes were centrifuged at 4,500 rpm for 1 min and 20 μl meiocytes suspension was transferred onto a *Polysine®* 2 *(Poly-L-Lysine coated slides)* and left to air dry. The immuno-cytology was done according to Colas *et al*. (2017). Briefly, the primary antibody solution consisted of anti-TaASY1 (rabbit), anti-HvZYP1 (rat) and anti-guinea pig HvDMC1 at 1:1000, 1:500 and 1:200, respectively in 1x PBS blocking buffer (Colas et al, 2017 & 2019). Primary antibodies were added on the slide, incubated in a wet chamber for 30 minutes at room temperature followed by up to 36 hours at 4 – 6°C. Slides were warmed for 1 hour at room temperature before washing for 15 minutes in 1x PBS and incubating for 90 minutes at room temperature in the secondary antibody solution consisting of a mixture of anti-rabbit Alexa Fluor® (488)with anti-rat Alexa Fluor® 568 and/or anti-guinea pig Alexa Fluor® 568(Invitrogen, Thermo Fisher Scientific) diluted in 1x PBS (1:600). Slides were washed for 15 minutes in 1x PBS and mounted in Vectashield® containing DAPI (H-2000, Vectorlabs). 3D Confocal stack images were acquired with LSM-Zeiss 710 using laser light (405, 488, 561 nm) sequentially. Stack images were processed using Imaris 8.1.2 (Bitplane).

### RNA extraction and sequencing

Total RNA was extracted from anthers, isolated meiocytes and EMB using TRIzol® PlusRNA Purification Kit (Thermo Fisher Scientific) following the manufacturer’s instructions. Total RNA was quality checked using a Bioanalyzer 2100 (Agilent). RNA-seq libraries were constructed using TruSeq mRNA Sample Preparation kit (Illumina) as recommended by the manufacturer. Libraries were quality checked using a Bioanalyzer 2100 (Agilent) and quantified by both qPCR (KAPA Library Quantification Kit, KAPA Biosystems) and using a Qubit fluorometer. RNA-seq was performed on a NextSeq 550 (Illumina) using recommended procedures, with all 6 anther and meiocyte libraries multiplexed on a single run (2x 75 bp, high-output). Total RNA samples from barley EMB were sequenced (Illumina PE150) by Novogene (HK) Company Limited, Hong Kong. Fastq files were used for downstream analysis.

Single molecule sequencing was performed at Earlham Institute, Norwich, UK using the PacBio Iso-seq method. Two RNA samples were prepared by mixing equal amounts of total RNA representing four or two different stages of isolated anthers or meiocytes, respectively. Anthers and meiocytes full-length cDNA libraries were prepared using 2 ug of total RNA and TeloPrime Full-length cDNA Amplification kit (TATAA Biocenter) according to the manufacturer’s instructions. PCR Optimisation was carried out on the two cDNA samples to determine the required number of cycles per individual sample. The final extension time was increased from 5 minutes to 7 minutes compared to the Teloprime method (Lexogen GmbH, Vienna, Austria) in order to ensure the generation of full-length PCR products, and each sample required between 16-18 PCR cycles respectively to generate sufficient cDNA for SMRTbell library preparation.

A third total RNA sample was extracted from a mixture of isolated anthers ranging from 0.3 to 1.2 mm in length. The corresponding full-length cDNA library was then generated using 1μg total RNA and the SMARTer PCR cDNA synthesis kit (Clontech, Takara Bio Inc., Shiga, Japan) following PacBio recommendations set out in the Iso-Seq method: https://www.pacb.com/wp-content/uploads/Procedure-Checklist-Iso-Seq-Template-Preparation-for-Sequel-Systems.pdf. PCR optimisation was carried out on the full-length cDNA using the KAPA HiFi PCR kit (Kapa Biosystems, Boston USA) and 12 cycles was sufficient to generate the material required for SMRTbell library preparation. The libraries were then completed following PacBio recommendations in the Iso-Seq method (see above link).

Each cDNA sample was bead cleaned with AMPure PB beads post PCR in preparation for SMRTbell library construction. SMRTbell library construction was completed following PacBio recommendations (above link). The cDNA libraries generated were quality checked using a Qubit Fluorometer 3.0 (Invitrogen) and sized using the Bioanalyzer HS DNA chip (Agilent Technologies, Inc.). The loading calculations for sequencing were completed using the PacBio SMRTlink Binding Calculator v5.0.0.6236. The sequencing primer from the SMRTbell Template Prep Kit 1.0-SPv3 was annealed to the adapter sequence of the libraries. Each library was bound to the sequencing polymerase with the Sequel Binding Kit v2.0 and the complex formed was then bound to Magbeads in preparation for sequencing using the MagBead Kit v2. Calculations for primer and polymerase binding ratios were kept at default values. Sequencing Control v2.0 was spiked into each library at ~1% prior to sequencing. The libraries were prepared for sequencing using the PacBio recommended instructions laid out in the binding calculator. The sequencing chemistry used to sequence all libraries was Sequel Sequencing Plate v2.1 and the Instrument Control Software version was v5.0.0.12545. The libraries were sequenced on the Sequel Instrument v1, using 1 SMRTcell v2.1 per library. All libraries had 600-minute movies, 120 minutes of immobilisation time, and 90 minutes pre-extension time.

### Barley Anther Transcriptome (BAnTr) construction

#### Illumina read mapping

The 12 anther samples, 6 meiocyte samples and 4 EMB samples of Illumina sequencing reads were mapped individually against the barley *cv.* Golden Promise assembly using STAR 2.7.1a (Dobin *et al.* 2013) in a two-step process based on Veeneman *et al.* (2016). The first genome index was generated using default settings. The exact parameters for the two-step mapping process can be found in Supplemental Methods.

#### PacBio read mapping

All three samples of long-read transcriptome data were processed individually using the SMRT Analysis software package v3.1.0 (https://github.com/PacificBiosciences/IsoSeq3) for building of the consensus circular reads (CCS) and demultiplexing which included barcode and primer removal. Afterwards all transcripts were merged for the clustering and polishing. Polishing on the merged transcripts was done using proovread v.2.14.1 (Hackl *et al.*, 2014) with the Illumina reads from the 12 anther samples and the 6 meiocyte samples. The polished transcripts were mapped to the Golden Promise reference assembly using GMAP version 2018-07-04 (Watanabe and Wu, 2005).

#### Transcriptome building

We used the Mikado pipeline (Mikado v1.2.4; Venturini *et al*., 2018) to join the different strands of transcriptome evidence, generated above. As additional transcript evidence from a more complete transcriptome we included the BaRTv1 transcripts (Rapazote-Flores *et al.*, 2019). The BaRTv1 transcripts were mapped to the Golden Promise reference assembly (GMAP version 2018-07-04; -n 0 --min-trimmed-coverage=0.80 --min-identity=0.90). Input files for the Mikado file included a splice junction file generated by Portcullis (Mapleson *et al*., 2017; default parameters), open reading frame identification of the transcripts by TransDecoder (https://github.com/TransDecoder/TransDecoder, default parameters; Haas *et al.*, 2013) and blast results using DIAMOND BLASTx (Buchfink *et al.*, 2015; --evalue 1e-5) against the NCBI non-redundant protein database as evidence. Together with the stringtie.gtf, scallop.gtf, pacbio.gtf and bartv1.gtf those were combined and scored, generating the Barley Anther Transcriptome (BAnTr). This dataset comprises 65,795 genes and 119,778 transcripts (Supplemental Dataset 6: AntherTranscriptomeBAnTr.fasta)

A padded version of the transcriptome was generated for the transcript quantification as previous experiments in *Arabidopsis* showed this to improve quantification and provide more similar results to high-resolution real-time PCRs (Zhang *et al.*, 2017; Supplemental Dataset 7: AntherTranscriptomeBAnTrPadded.fasta).

To distinguish between protein-coding and long non-coding transcripts, all transcripts were run through the NCBI ORF finder. Transcripts and corresponding genes were assigned as coding, if one hit was identified in a DIAMOND BLASTp (v0.9.24; Buchfink *et al.*, 2015) search [--query-cover 60 --evalue 1e-30] against the NCBI non-redundant protein database (Supplemental Dataset 8: AntherProteomeBAnTr.fasta). Additionally, transcripts were checked using CPC2 (coding potential calculator 2, run on both strands) and InterProScan (v5.39, default parameters; Jones *et al.*, 2014). If both supported a coding sequence, those were also included in the coding transcripts; if only one of those provided evidence for coding, the sequences were allocated to the unclassified group. All remaining transcripts and their corresponding genes were assigned as long non-coding RNA.

### Differential expression

To study differential gene expression the 12 anther and 6 meiocyte Illumina sequencing samples were mapped against our above described padded BAnTr transcriptome using Salmon v.0.14.1 (Patro *et al.*, 2017; -validateMappings --useVBOpt --seqBias --gcBias --posBias). The subsequent analysis was conducted using the 3D RNA-seq pipeline (Guo *et al.*, 2019; accessed 28/11/2019). This pipeline is a combination of different R packages. Transcript abundance is imported using the lengthScaledTPM method from tximport (Soneson *et al.*, 2016) and transformed into gene level counts or kept as transcript level counts. Low expressed transcripts with a count per million reads (CPM) below 1 in less than three samples were removed. This reduced the dataset to 31,918 genes and 50,861 transcripts. Gene and transcript read counts were normalised using the trimmed mean of M values (TMM) method, implemented in the edgeR package (Robinson *et al.*, 2010; Robinson and Oshlack 2010; McCarthy *et al.*, 2012). Based on the experimental design, six contrast groups were built: anthers at leptotene–zygotene versus anthers at pre-meiosis; anthers at pachytene–diplotene versus anthers at leptotene–zygotene; anthers at metaphaseI–tetrad versus anthers at pachytene–diplotene; meiocytes at leptotene–zygotene versus anthers at leptotene–zygotene; meiocytes at pachytene–diplotene versus anthers at pachytene–diplotene; and meiocytes at pachytene–diplotene versus meiocytes at leptotene–zygotene. Differential expression analysis was done using the voom function in the limma R package (Law *et al.*, 2014; Ritchie *et al.*, 2015). Genes and transcripts were rated as differentially expressed with a log_2_ fold change (log_2_FC) above 1 or below −1 and an adjusted p-value (by the Benjamini-Hochberg method (Benjamini and Hochberg, 1995)) of below 0.01.

#### Differential alternative splicing analysis

Differential alternative splicing analysis is part of the 3D RNA-seq pipeline. It integrates the diffSplice function from the limma R package and compares the log_2_FC of each individual transcript with the gene level log_2_FC. Transcripts with a Δ percent spliced (ΔPS) ratio of above 0.1 were assigned as significant alternative spliced transcripts. To determine the alternative splicing events, we used SUPPA version 2.3 (Alamancos *et al.*, 2015; Trincado *et al.*, 2018).

#### Co-expression analysis

To identify co-expressed networks, we used the weighted gene co-expression network analysis (WGCNA) R package (v1.63; Langfelder and Horvath, 2008). The same reduced dataset of 31,918 expressed genes as introduced above was used as input. For the network construction an approximate scale-free topology of above 0.85 was achieved with a soft power of 10. Network construction and module assignment was done with the following settings: a signed hybrid network, mergeCutHeight of 0.35, a minModuleSize of 30 and a deepSplit of 4.

##### Gene Ontology and functional annotation

Multiple strands of evidence were combined for the functional annotation of the proteins. Mercator4 v.2 (Schwacke *et al.*, 2019) was used to classify the proteins into functional bins. eggNOG (Huerta-Cepas *et al.*, 2017; Huerta-Cepas *et al.*, 2019) was used to add COG (Clusters of Orthologous Groups of proteins) annotation and a protein description. PANNZER2 (Toronen *et al.*, 2018) was used for the GO term identification and a protein description. The GO enrichment analysis was done using the topGO R package using the fisher weight01 algorithm and a P value cut-off below 0.001 (Alexa *et al.*, 2006). TopGO only outputs an exact P value until 1e-30. Everything below will be given as P < 1e-30. For visual presentation in the GOplot R package (Walter *et al.*, 2015) those were set to 1e-30.

## Accession numbers

The raw data is available from the NCBI-Sequence Read Archive database. Illumina sequences from anthers and meiocytes samples and the PacBio sequences can be found in the Bioproject PRJNA558196. Illumina sequences of four EMB samples can be found in the Bioproject PRJNA593943.

## Acknowledgements

This work was funded by the European Research Council (ERC Shuffle Project ID: 669182) and supported by the Scottish Government’s Rural and Environment Science and Analytical Services Division work programme Theme 2 WP2.1 RD1 and RD2. We would like to thank Dr Paulo Rapazote-Flores and Dr Runxuan Zhang for their Bioinformatic help and suggestions and Mr. Philip Smith for proof reading.

## Author contributions

AB, DL, IC, MS, LR and RW, experimental design; NM, plant growth; AB, DL, IC, MA, MM, MAS and NM, collection of anthers; IC, collection of meiocytes and immunocytology; AB, PH and JM, RNA extraction, library prep, QC; MS, transcriptome assembly; AB, lead manuscript preparation; AB, DL, JO, IC, MS and RW, data analysis and manuscript preparation.

## SUPPLEMENTAL FILES

### SUPPLEMENTAL FIGURES

**Supplemental Figure 1:** Sample clustering analysis by 3D multidimensional scaling plot.

A.PRE, anther pre-meiosis; A.LEP-ZYG, anther leptotene–zygotene; A.PAC-DIP, anther pachytene–diplotene; A.MET-TET, anther metaphase I–tetrad; M.LEP-ZYG, meiocyte leptotene–zygotene; M.PAC-DIP, meiocyte pachytene–diplotene.

**Supplemental Figure 2:** Overview from the 3D RNA-seq results.

**a)** Differential expressed genes and differential alternative spliced genes. **b)** Differential expressed transcripts and differential transcript usage transcripts. DE, differential expressed; DAS, differential alternative spliced; DTU, differential transcript usage; AS, alternative splicing

**Supplemental Figure 3:** WGCNA analysis of co-expressed genes.

A total of 17 modules were found in anther and meiocyte transcriptomes.

**Supplemental Figure 4:** Differential eigengene network analysis.

**a)** Clustering dendrogram of consensus module eigengenes. **b)** Heatmap of eigengene adjacencies in barley anther and meiocyte samples. Each row and column correspond to one eigengene labelled by consensus module colour in the band. Within the heat map black indicates high adjacency (positive correlation) and white low adjacency (no correlation) as shown by the colour legend.

**Supplemental Figure 5:** Gene ontology enrichment of the red module of the WGCNA.

Size of bubbles correspond to total number of proteins associated with the GO term. MF = molecular function, BP = biological process, CC = cellular component.

**Supplemental Figure 6:** Heatmap of meiotic gene expression.

Only genes without a statistically significant log fold change in expression are shown. Genes are ordered and labelled by WGCNA module on the vertical axis. The samples (3 replicates each) are A.PRE, anther pre-meiosis; A.LEP-ZYG, anther leptotene–zygotene; A.PAC-DIP, anther pachytene–diplotene; A.MET-TET, anther metaphase I–tetrad; M.LEP-ZYG, meiocyte leptotene–zygotene; M.PAC-DIP, meiocyte pachytene–diplotene.

**Supplemental Figure 7:** A heatmap of transcript levels of genes involved in post-transcriptional regulation.

Changes of expression levels of different genes involved in small RNA pathways and N 6-methyladenosine (m^6^A) mRNA methylation were determined by comparing meiocytes and anthers of the same developmental stage or anthers only at different stages. Transcript counts for each gene were extracted from total expression data, log transformed to normalise distribution, and plotted using ggplot2 (Wickham, 2016) in R (this study, available at https://github.com/BioJNO/BAnTr). The samples (3 replicates each) are A.PRE, anther pre-meiosis; A.LEP-ZYG, anther leptotene–zygotene; A.PAC-DIP, anther pachytene–diplotene; A.MET-TET, anther metaphase I–tetrad; M.LEP-ZYG, meiocyte leptotene–zygotene; M.PAC-DIP, meiocyte pachytene–diplotene. AGO: argonaute gene family.

### SUPPLEMENTAL TABLES

**Supplemental Table 1**: Number of genes per module based on the WGCNA.

**Supplemental Table 2:** Differential expressed genes and differential expressed transcripts

### SUPPLEMENTAL DATA SETS

**Supplemental Data set 1:** GO counts in different WGCNA modules.

**Supplemental Data set 2:** GO counts in different comparisons.

**Supplemental Data set 3:** Meiocyte enriched ligases.

**Supplemental Data set 4:** Differential expressed transcription factor families.

**Supplemental Data set 5:** Distribution of transcription factor families in different modules.

**Supplemental Data set 6:** AntherTranscriptomeBAnTr.fasta.

**Supplemental Data set 7:** AntherTranscriptomeBAnTrPadded.fasta.

**Supplemental Dataset 8**: AntherProteomeBAnTr.fasta.

**Supplemental Video 1:** Isolated fresh meiocytes bag at Leptotene/zygotene.

**Supplemental Video 2:** Isolated fresh meiocytes bag at pachytene/diplotene.

